# Defining *Escherichia coli* as a health-promoting microbe against intestinal *Pseudomonas aeruginosa*

**DOI:** 10.1101/612606

**Authors:** Theodoulakis Christofi, Stavria Panayidou, Irini Dieronitou, Christina Michael, Yiorgos Apidianakis

## Abstract

Gut microbiota acts as a barrier against intestinal pathogens, but species-specific protection of the host from infection remains relatively unexplored. Taking a Koch’s postulates approach in reverse to define health-promoting microbes we find that *Escherichia coli* naturally colonizes the gut of healthy mice, but it is depleted from the gut of antibiotic-treated mice, which become susceptible to intestinal colonization by *Pseudomonas aeruginosa* and concomitant mortality. Reintroduction of fecal bacteria and *E. coli* establishes a high titer of *E. coli* in the host intestine and increases defence against *P. aeruginosa* colonization and mortality. Moreover, diet is relevant in this process because high sugars or dietary fat favours *E. coli* fermentation to lactic acid and *P. aeruginosa* growth inhibition. To the contrary, low sugars allow *P. aeruginosa* to produce the oxidative agent pyocyanin that inhibits *E. coli* growth. Our results provide an explanation as to why *P. aeruginosa* doesn’t commonly infect the human gut, despite being a formidable microbe in lung and wound infections.

**Author Summary:** Here we interrogate the conundrum as to why *Pseudomonas aeruginosa* is not a clinical problem in the intestine as opposed to other tissues. *P. aeruginosa* interacts with Neisseria, Streptococcus, Staphylococcus and Actinomyces species found in the human lung. These are predominantly gram-positive bacteria that induce *P. aeruginosa* virulence. Moreover, peptidoglycan, which is abundant in gram-positive bacteria, can directly trigger the virulence of *P. aeriginosa*. We reasoned that *P. aeruginosa* might be benign in the human gut due to the inhibitory action of benign gram-negative intestinal bacteria, such as *Escherichia coli*. Therefore, we dissected the antagonism between *E. coli* and *P. aeruginosa* and the effect of a conventional, a fat-, a carbohydrate-and a protein-based diet in intestinal dysbiosis. Our findings support the notion that an unbalanced diet or antibiotics induces gut dysbiosis by the elimination of commensal *E. coli*, in addition to lactic acid bacteria, imposing a gut environment conducive to *P. aeruginosa* infection. Moreover, commensal *E. coli* provides an explanation as to why *P. aeruginosa* doesn’t commonly infect the human gut, despite being a formidable microbe in lung and wound infections.

## Introduction

*Escherichia coli* and *Streptococci* are the first bacteria to colonize the gastrointestinal tract of humans following birth and seem to shape the environment in the gut for the establishment of other bacteria such as *Bifidobacterium* and *Bacteroides* (Mitsuoka 1996, Mackie et al. 1999). *Bifidobacterium* and *Lactobacillus* strains are considered the main fermenters in the human gut (Zoetendal et al. 2012, Macfarlane, Macfarlane 2011). *E. coli* thrives aerobically, but also in the anaerobic mammalian gut where it ferments carbon sources to produce short chain fatty acids and other metabolic by-products (Mitsuoka 1996, Winter 2013). Moreover, gut inflammation generates host-derived nitrate that confer a fitness advantage to commensal *E. coli* (Winter et al. 2013). Interestingly, the probiotic *E. coli* Nissle 1917 (EcN) is particularly beneficial for ulcerative colitis patients in maintaining disease remission (Kruis et al. 1997, Kruis et al. 2004, Matthes et al. 2010). In addition, EcN induces host immune defence against pathogens (Boudeau et al. 2003, Schlee et al. 2007) strengthens intestinal barrier (Zyrek et al. 2007, Ukena et al. 2007) and directly inhibits pathogenic *E. coli* strains (Maltby et al. 2013, Reissbrodt et al. 2009). Yet, the beneficial role of *E. coli* is so far true only for EcN, while lactic acid bacteria, such as Lactobacilli and Bifidobacteria, are considered the main fermenters in the mammalian gut that may directly inhibit pathogens.

Antibiotics can also greatly affect microbiota diversity and promote dysbiosis. Early in life, antibiotic treatment can modulate the development of gut microbiota in children (Tanaka et al. 2009). In children and adults, opportunistic pathogens can take advantage of the antibiotic effect on commensal bacteria to infect the gut (Chang et al. 2008). One such pathogen is the gram-negative human opportunistic bacterium *Pseudomonas aeruginosa*, one of the most frequent species in hospital-acquired infections (Markou and Apidianakis 2013). While not a common clinical problem in the gut, *P. aeruginosa* colonizes the gastrointestinal tract of many hospitalized patients and to a lesser extent of healthy individuals (Ohara and Itoh 2003, Shimizu et al. 2006, Vincent et al. 2009, Chuang et al. 2014). *P. aeruginosa* can nevertheless cause frequent and sever wound and lung infections in immunocompromized individuals, and the ears and eyes of seemingly healthy people (Panayidou and Apidianakis 2017). It is responsible for more than 50,000 infections per year in the U.S., causing acute, chronic and relapsing/persistent infections due to a wide variety of virulence factors. Many of its virulence genes are controlled by quorum sensing (QS), a communication system that promotes synchronized bacterial behaviors, such as the production of the oxidative agent pyocyanin (Lau et al. 2004).

Here we interrogate the contribution of *E. coli* in controlling *P. aeruginosa* intestinal colonization in a diet-dependent manner. We apply the Koch’s postulates in reverse to prove causation of commensal *E. coli* in fending off *P. aeruginosa* infection. Accordingly we find that: (**a**) a mouse *E. coli* strain is detected through culture-independent methods (16S sequencing) in the feces of untreated, but not of antibiotically-treated mice, which are susceptible to infection; (**b**) Candidate health-promoting commensal *E. coli* strain species was isolated through culture-dependent microbiological analysis and archived as a pure culture in the laboratory; (**c**) This mouse *E. coli* and other *E. coli* strains ameliorate *P. aeruginosa* infection when introduced into antibiotic-treated mice; (**d**) The administered health-promoting *E. coli* strains can be identified in high titers in the feces of mice to which resistance to infection was improved. Moreover, assessing three extremes and a conventional diet in mice we find that, while sugar is fermented by various *E. coli* strains to lactic acid in culture, in the mouse gut a vegetable fat-based rather than a carbohydrate-or protein-based diet boosts lactic acid production and helps *E. coli* to inhibit *P. aeruginosa*. Our findings support the notion that unbalanced diets or the use of antibiotics may eliminate not only lactic acid bacteria but also commensal *E. coli*, imposing a gut environment conducive to *P. aeruginosa* infection.

## Methods

### Bacterial stains

*Pseudomonas aeruginosa* strain UCCBP 14 (PA14) and isogenic gene deletion mutants *Δmvfr, ΔphzS, ΔphzS* and *ΔrhlR/ΔlasR* are previously described (Rahme et al. 1995; Kapsetaki et al. 2014). *E. coli* MGH and *E. coli* BWH and all other strains used (but Lactobacillus ones) are human isolates obtained from Prof. Elizabeth Hohmann at Mass General Hospital and Prof. Andrew Onderdonk at Brigham and Women’s Hospital. Lactobacillus strains are isolated from wild-caught *Drosophila* and are previously described (Chandler et al. 2011). Mouse *E. coli (E. coli* CD1) was isolated from feces of CD1 mice for this study and validated through colony PCR and biochemical analysis [positive Indole production and positive growth on selective chromogenic Tryptone Bile X-glucuronide (TBX) agar plate]. Laboratory *E. coli* BW25113 and KEIO collection strains, including *Δpgi, ΔadhE, ΔatpC, Δpta* and *ΔldhA* are previously described (Baba et al. 2006). Laboratory *E. coli* BW25113 and *Δtna, ΔsdiA, ΔluxS,* strains are previously described (Chu et al. 2012). Enteropathogenic (EPEC) *E. coli* O127:H6 E2348/69 was obtained from Prof. Tassos Economou and was previously described (Levine et al. 1978).

### Bacteria handling for in culture experiments

*E. coli* and *P. aeruginosa* strains were grown at 37°C overnight with shaking at 200 rpm in liquid LB from frozen LB-20% glycerol stocks cultures and then were diluted to OD_600nm_ 0.01 in fresh sterile LB to establish mono-or co-cultures. Sucrose or glucose was added to a final concentration of 4% w/v during growth assessments. Bacterial supernatants were produced by overnight bacterial cultures filter-sterilized and mixed in 1:1 volume ratio with fresh LB broth. Selective plates contained 50μg/ml rifampicin for *P. aeruginosa* and 60 μg/ml kanamycin for *E. coli* Keio collection or TBX agar for wild type *E. coli*.

### Fly survival

Aerobic strains were grown at 37°C overnight with shaking at 200 rpm in liquid LB from frozen LB-20% glycerol stocks cultures and then diluted to OD_600nm_ 0.01 in fresh sterile LB grown over day to OD_600nm_ 3. Anaerobic strains were grown at 37°C for 72 hours without shaking in liquid BHI from frozen BHI-20% glycerol stocks to OD_600nm_ 1-2. Cultures were then pelleted and diluted to a final OD_600nm_ 0.15 per strain in a 4% sugar (sucrose or glucose), 10% sterile LB infection medium. Wild type Oregon R *Drosophila melanogaster* female flies 3-5 days old were starved for 6 hours prior to infection. 5ml infection medium was added on a cotton ball at a bottom of a fly vial. Each vial contained 10 to 15 flies and observed twice a day for fly survival (Apidianakis et al 2009).

### Fly colonisation

Germ free flies are generated through dechorionation of collected eggs in 50% bleach. Adult Oregon R 3-5 day old flies were infected for 24 hours with a mix of bacterial culture(s) grown as mentioned above, pelleted and diluted to a final OD_600nm_ 0.02 per strain in a 4% sugar (sucrose or glucose) medium. Flies were then transferred and maintained in modified falcon tubes with 200μl 2% or 4% of sucrose or glucose as previously described (Kapsetaki et al. 2014). At day 2 and day 5 flies were homogenised using the Qiagen Tissuelyser LT for 5 minutes at 50Hz. Bacteria CFUs were enumerated on selective plates after overnight incubation at 37°C.

### KEIO *E. coli* gene deletion library screen

The Keio *E. coli* collection of gene knockouts was acquired from the Japanese National Institute of Genetics and contains 3884 *E. coli* mutants with unique gene deletions. Strains were grown overnight in sterile 96 well clear flat bottom plates containing 200 μl of sterile LB broth at 37°C and 200rpm shaking. *P. aeruginosa* was grown in glass tubes as standard overnight conditions. Over day co-cultures were performed at 37oC and 200rpm in 96 well plates containing 1:100 *P. aeruginosa* and 1:100 *E. coli* mutant overnight inoculations in 200μl LB Broth supplemented with 4% glucose. At 24 hours Pyocyanin production was observed visually using as positive controls PA14 monocultures and co-cultures of PA14 with *E. coli* mutants lacking inhibitory properties (e.g. *Δpgi*). Bacterial growth was measured at OD_600nm_ on a plate reader. Bacterial co-cultures typically exhibit half the optical density of PA14 monocultures. Thus co-cultures with equal or higher the optical density of PA14 monocultures indicated antagonistic interactions.

### Animal Diets

*Drosophila melanogaster* Oregon R flies were reared in cornmeal, yeast and sugar diet at 25°C in a 12 hour day and night cycle. CD1 mice were reared 5-6 individuals per cage at 24°C in a 12 hour day and night cycle. Standard chow diet was obtained from Mucedola s.r.l Italy (#4RF25 a complete balanced diet containing mainly Starch 35.18%, Sucrose 5.66%, Crude Protein 22%, Crude oil 3.5%). Specialised diets based on either vegetable fats, carbohydrates or protein were manufactured by Mucedola s.r.l (#PF4550, PF4551 and PF4552) per Table 1 below (Smith et al. 1999).

**Table 1.** Composition of macronutrient diets (% by weight)

|  | Carbohydrate | Fat | Protein |
| --- | --- | --- | --- |
| Cornstarch | 58.11 | 0.00 | 0.00 |
| Powdered sugar | 29.06 | 0.00 | 0.00 |
| Casein | 0.00 | 0.00 | 87.17 |
| dl-Methionine | 0.11 | 0.20 | 0.11 |
| Vegetable shortening* | 0.00 | 75.12 | 0.00 |
| AIN-76A vitamin mix** | 0.77 | 1.49 | 0.77 |
| AIN-76A mineral mix** | 3.07 | 5.95 | 3.07 |
| Choline chloride | 0.18 | 0.34 | 0.18 |

**Table 1. Composition of macronutrient diets (% by weight)**
|  | Carbohydrate | Fat | Protein |
| --- | --- | --- | --- |
| Cellulose (Alphacel) | 8.72 | 16.91 | 8.72 |
| Energy density, kcal/g | 3.53 | 6.85 | 3.53 |

\* Crisco brand, a blend of soybean oil, fully hydrogenated palm oil, and partially hydrogenated palm and soybean oils. Contains 50% polyunsaturated fat, 20.8% monounsaturated fat, 0% trans fat and 25% saturated fat per weight.
\*\*Vitamin (A and D3) and mineral (Fe, Mn, Zn, Cu, I, Se) mixes contain 97% and 12% sucrose, respectively.

### Ethics Statement

Animal protocols have been approved by the Cyprus Veterinary Service inspectors under the license number CY/EXP/PR.L6/2018 towards the Laboratory of Prof. Apidianakis at the University of Cyprus. The veterinary services act under the auspices of the Ministry of Agriculture in Cyprus and the project number CY.EXP101. These national services abide to the National Law for Animal Welfare of 1994 and 2013 and the Law for experiments with animals of 2013 and 2017.

### Mouse colonisation assay

Female CD1 mice 7-8 weeks old are treated with an antibiotic cocktail of 0.1mg/ml Rifampicin, 0.3 mg/ml Ampicillin and 2 mg/ml Streptomycin for 6 days to reduce endogenous gut bacteria. Starting the next day PA14 was provided daily for 7 days in the drinking water prepared from an over day culture of OD_600nm_ 3, centrifuged at 7000 rpm (4610 RCF) for 5 minutes to collect bacteria and diluted 1:10 to obtain ∼3×10^8^ bacteria/ml. Following infection (Day 0 PA14 colonisation) *E. coli* was provided for 1 day at the same concentration and CFUs for both bacteria were measured every other day from homogenized and plated mouse feces.

### 16S Metagenomic

Mouse fecal samples were collected in eppendorf tubes, weighed, snap frozen and stored at −80°C. Bacterial DNA was extracted using the QIAamp DNA Stool Mini Kit. 16S Sequencing was performed by the Illumina metagenomics analyser. Kraken software was used to assign taxonomic sequence classification.

### Mouse survival assay

Female CD1 mice 7-8 weeks old were supplied in drinking water with an antibiotic cocktail of 0.1mg/ml Rifampicin, 0.3 mg/ml Ampicillin and 2 mg/ml Streptomycin for 6 days to reduce endogenous gut bacteria. Starting the next day culture of specific *E. coli* strains was provided in drinking water for 24 hours prepared from an over day culture of OD_600nm_ 3 and/or anaerobic fecal culture grown to the maximum (2 Days) centrifuged at 7000 rpm (4610 RCF) for 5 minutes to collect bacteria and diluted 1:10 to obtain ∼3×10^8^ bacteria/ml. The next day P. aeruginosa (strain PA14) was provided daily for 7 days in the drinking water as for *E. coli*. Then mice were injected intraperitoneally with 150mg/kg of body weight with cyclophosphamide and 3 days later with another dose of 100mg/kg as previously described (Zuluaga et al. 2006). Survival was observed twice a day until all mice die or for up to 1 week.

### Acid and Sugar measurements

Lactic and acetic acid concentrations in culture supernatants and homogenised mouse feces (produced via bead homogenization in water) were determined enzymatically using R-Biopharm kits No. 11112821035 and No. 10148261035 respectively, according to manufacturer’s instructions. Sugar concentrations in homogenised mouse feces were determined using the Megazyme Sucrose/D-Fructose/D-Glucose Assay Kit (K-SUFRG) according to manufacturer’s instructions. Absorbance was measured using the NanoDrop 2000c Spectrophotometer.

### Pyocyanin measurement

Overnight PA14 cultures were diluted to OD_600nm_ 1, then 0.25ml was used to inoculate 25ml of LB. Cultures were grown at 37°C, 200rpm in 250 ml flasks. Supernatants were collected after centrifugation at 6000 rpm (4800 RCF) for 10 minutes. 4.5 ml of chloroform was added to 7.5 ml of supernatant and vortexed. Samples were then centrifuged at 6000 rpm (4800 RCF) for 10 minutes. 3 ml of the resulting blue layer at the bottom was transferred to a new tube. 1.5 ml of 0.2 M HCl was added to each tube and vortexed 2 times for 10 seconds. Samples were centrifuged for 3 minutes at 6000 rpm (4800 RCF) and 1 ml of the pink layer was transferred to cuvettes. Pyocyanin concentration (μl/ml) was calculated by multiplying the spectrophotometric measurements taken at 520 nm by 17.072, then multiplying it again by 1.5 due to the chloroform dilution.

### Computational analysis

Pairwise comparison of bacterial CFUs, pH3 positive cells and other measurements were evaluated using the two-sided Student’s t-test for samples of ≥10 or U-test for samples <10. Survival curves of mice and flies were analyzed with the Kaplan-Meier method and the log-rank test. All experiments were repeated at least twice with qualitatively similar results. Gene enrichment analysis was performed using the David’s functional annotation tool. Correlation coefficient (*r*) significance analyses of mouse fecal acid concentration vs. LT50 was done using Pearson correlation and an n=6 (the average of six dietary conditions sampling 6 mice for each). Acetic + Lactic acid Index for each of the 6 dietary conditions was computed by dividing each acid concentration of each dietary condition with the average concentration of that acid in all conditions and adding the normalized values of the two acids. For sucrose assimilation prediction we used BLASTN 2.8.1+ per Zhang et al. 2000 we found (a) an *E. coli W* sucrose hydrolase (98% identity), (b) a sucrose permease (98% identity), (c) a sucrose-specific IIBC component (100% identity) and (d) a sucrose-6-phosphate hydrolase (100% identity) present in *E. coli* O127:H6 str. E2348/69 (taxid:574521), but not in the genomes of *E. coli* BW25113 (taxid:679895) and *E. coli* DH5[alpha] (taxid:668369).

## RESULTS

### *Escherichia coli* secreted factors antagonise *Pseudomonas aeruginosa* growth in the presence of sugars

Through a bacterial interaction screen we found that all flies orally infected with *P. aeruginosa* died within 6 days, but none of the *E. coli* strains tested (MGH, EPEC, BW25113 and DH5a) were that lethal to flies (LT50%>10 days; Figures 1A, S1A). However, we noticed a prominent increase in survival of *P. aeruginosa* infected flies when co-infected with either the human isolate *E. coli* MGH (Figure 1A) or the laboratory strain *E. coli* BW25113 (Figure S1A) in the presence of 4% sucrose or 4% glucose, respectively. Noticeably, sucrose can be used by *E. coli* MGH and EPEC to inhibit *P. aeruginosa* (LT50%>10 days; Figure 1), because EPEC, for example, possesses 4 sucrose uptake and metabolism enzymes [an *E. coli* W sucrose hydrolase (98% identity), a sucrose permease (98% identity), a sucrose-specific IIBC component (100% identity) and a sucrose-6-phosphate hydrolase (100% identity)]. To the contrary, the *E. coli* strains BW25113 and DH5a do not have these genes and were unable to utilize sucrose to inhibit *P. aeruginosa* in our experiments (LT50%<7 days). Nevertheless, when 4% glucose was used instead of sucrose in the infection mix, *E. coli* BW25113 gained the capacity to inhibit *P. aeruginosa* (Figure S1) (Jahreis et al. 2002).

**Figure 1:**
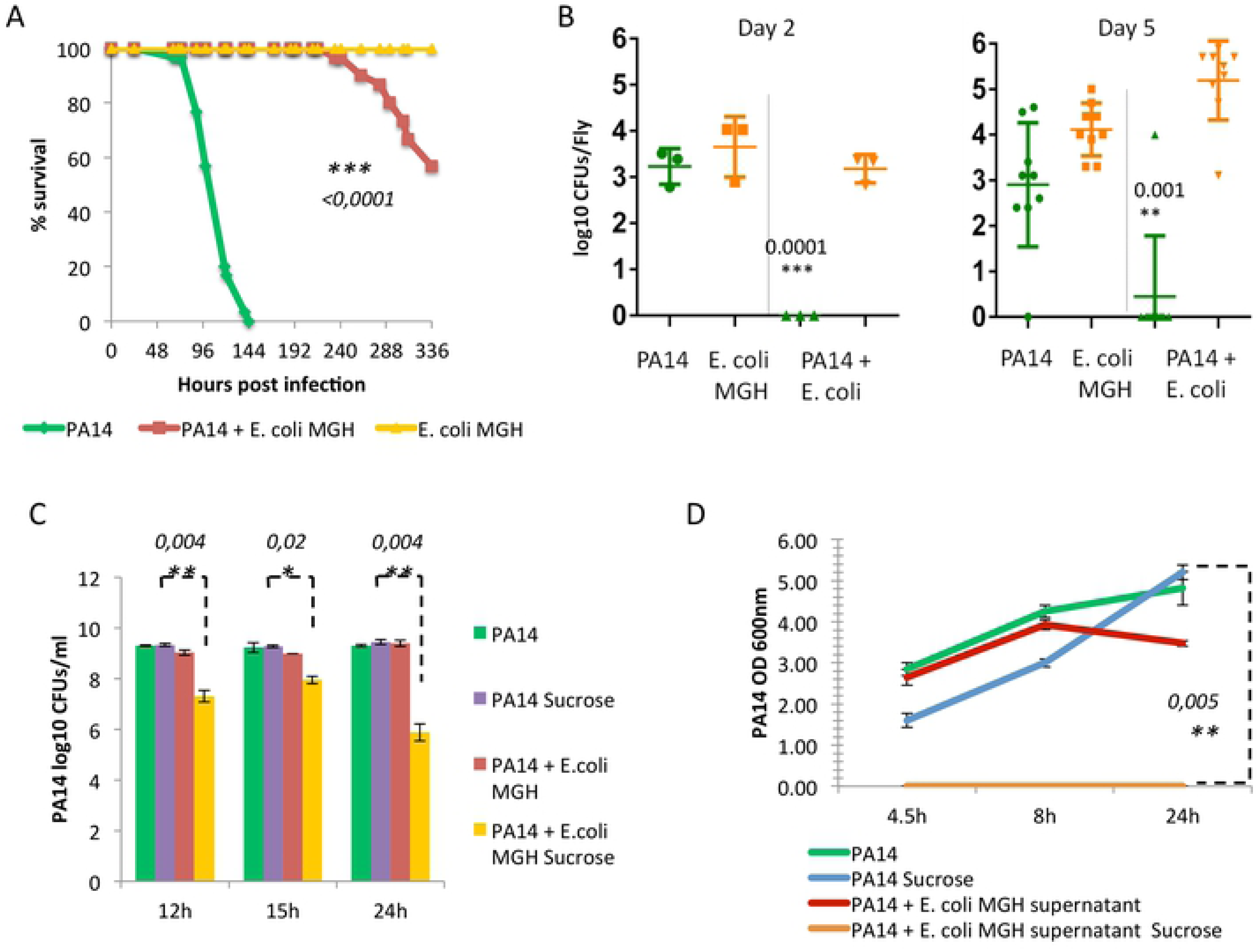
E. coli inhibits P. aeruginosa growth and virulence in the Drosophila gut and in culture in the presence of sucrose. A: Survival of Drosophila melanogaster Oregon R flies infected with PA14 (green), E. coli strain MGH (yellow) or co-infected with E. coli MGH and P. aeruginosa PA14 (red) [n=45]. B: Colonisation levels measured in colony forming units (CFUs) at Day 2 or Day 5 post PA14 infection (green), MGH (yellow) in mono or co-infected flies [n=3 and n=9]. C: CFUs of PA14 growth in the presents or absence of 4% sucrose and E. coli MGH in Luria-Bertani cultures [n=3-6]. D: Optical density measurements at 600nm of PA14 growth in liquid supernatant of E. coli cultures +/- 4% sucrose [n=9].

To assess if *E. coli* antagonizes *P. aeruginosa* by inhibiting its colonization, we assessed the bacterial loads of each bacterium in the fly upon co-infection. Colony forming unit (CFU) measurements in selective media revealed that at 2 days and 5 days after infection with either *P. aeruginosa* or *E. coli* MGH flies harboured roughly 10^3^ bacteria or more per fly (Figure 1B). Upon co-infection with *P. aeruginosa* and *E. coli* MGH, *P. aeruginosa* was almost eradicated, while *E. coli* MGH remained stable (Figure 1B). Using the *E. coli* BW25113 strain and *P. aeruginosa* to co-infect flies no significant changes were found in CFUs at day 2, but at day 5 of co-infection CFUs for both species were found tentatively lower, suggesting mutual inhibition at the level of colonization (Figure S1B).

To assess if the interaction between *P. aeruginosa* and *E. coli* is direct we assessed bacterial growth in LB culture. Interestingly, *E. coli* MGH did not inhibit *P. aeruginosa* growth in plain liquid LB (Figure 1C). To assess if sucrose added in the fly infection media as the main carbon source for the flies plays a role in bacterial interaction, we supplemented LB media with 4% sucrose. Strikingly, in the presence of sucrose *P. aeruginosa* concentration was reduced by >1,000 fold when co-cultured with *E. coli* MGH, but no inhibition was noticed in the absence of added sucrose (Figure 1C). The monosaccharides glucose and fructose enable also *E. coli* BW25113 to inhibit *P. aeruginosa* (Figure S1C). To assess whether secreted factors are responsible for *P. aeruginosa* growth inhibition we grew *P. aeruginosa* in a mix of 50% fresh LB and 50% filtered LB supernatant from an overnight *E. coli* culture that was supplemented or not with 4% of sugars. The mix containing supernatant of *E. coli* MGH grown in sucrose and that of *E. coli* BW25113 grown in glucose was able to completely inhibit the growth of *P. aeruginosa* for at least 24 hours (Figure 1D, S1D).

### *E. coli* inhibits *P. aeruginosa* intestinal colonization and lethality during mouse gut-derived sepsis

To model antibiotic induces dysbiosis we used a mouse assay of intestinal infection. We administered a regime of three broad-spectrum antibiotics in mice and assessed their gut microbiota at the genus level through 16S sequencing analysis. In the absence of antibiotics the microbiota consisted primarily of *Bacteroidetes, Firmicutes* and *Proteobacteria,* including *E. coli* (Figure 2A). Using colony PCR sequencing we verified the presence of *E. coli* (named the strain CD1) and identified 7 easy to culture and potentially beneficial strains belonging to the *Lactobaccillus, Bifidobacterium* and *Bacteroides* genera in the feces of mice (Figure 2H). Antibiotic treatment induced dysbiosis, which is exemplified by the eradication of *E. coli*, the reduction of all the prevalent phyla below the detection level (Figure 2B), and the eradication of all 8 cultured bacterial strains (Figure 2H), but *Bifidobacterium sp.2*, which was reduced from 8.4 log_10_ to 7.2 log_10_ CFUs per gram of mouse faeces.

**Figure 2:**
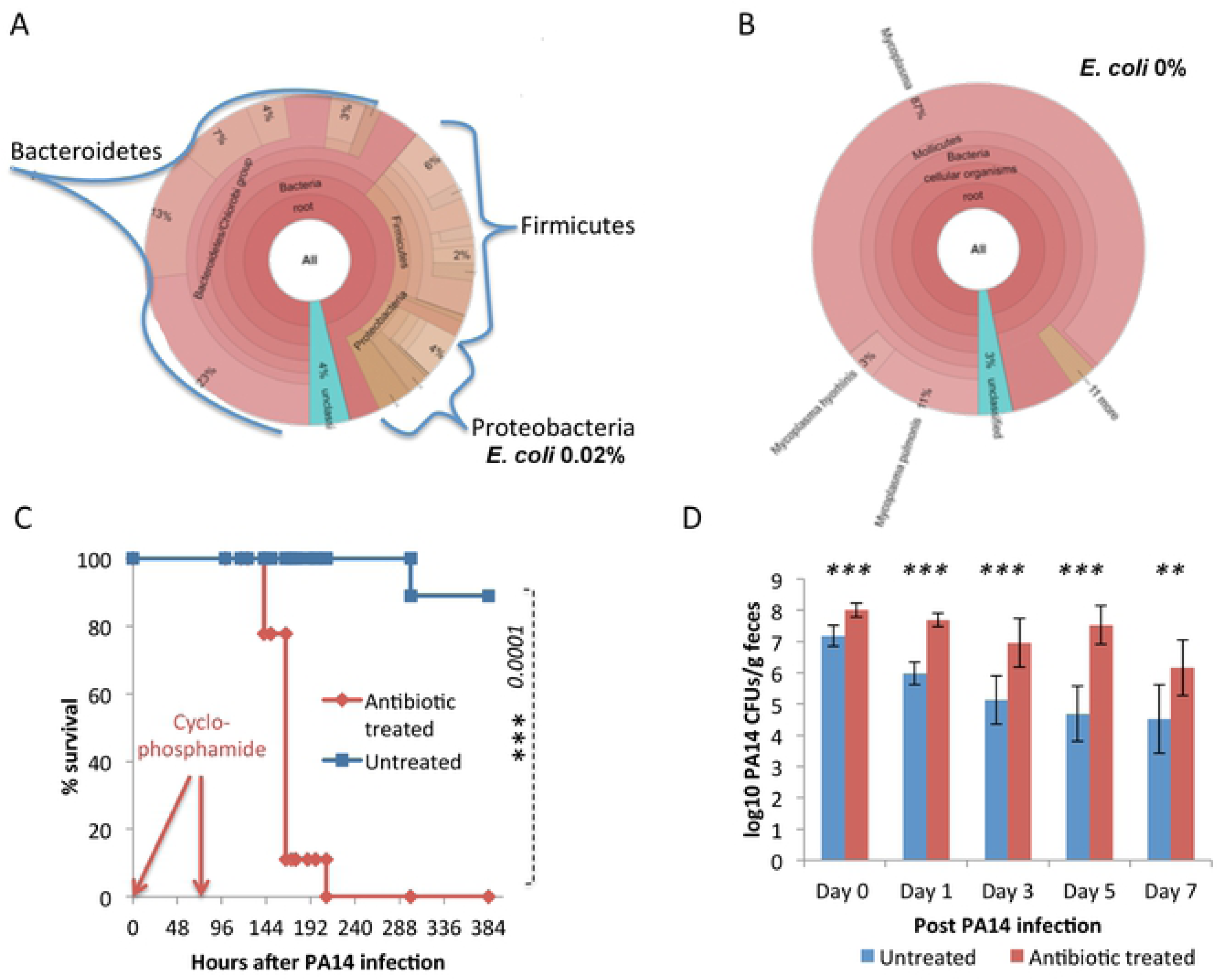
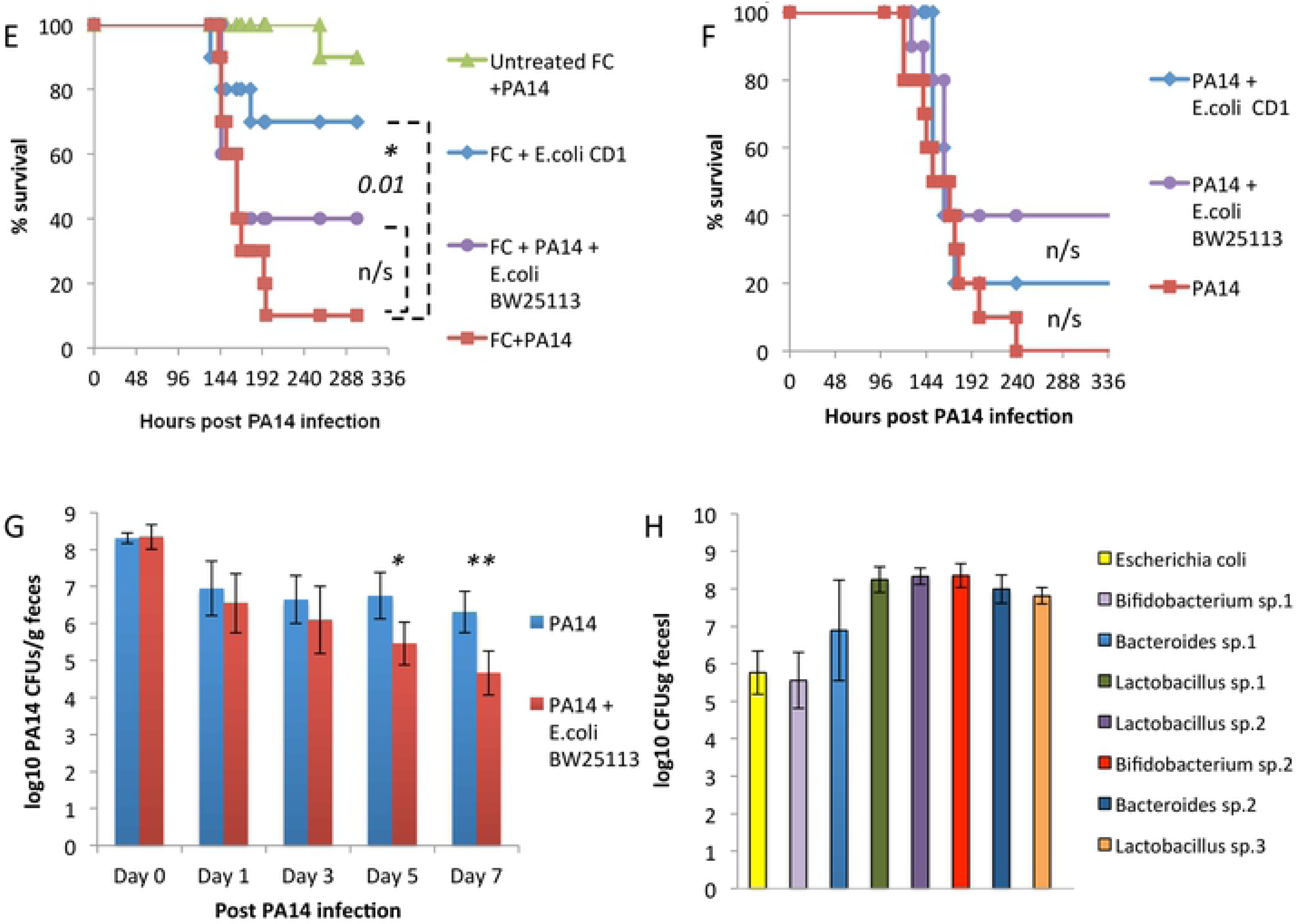
The use of antibiotics and protection of commensal bacteria (E. coli) against P. aeruginosa-induced lethality and colonization. A: 16S Metagenomics analysis of mouse feces before or B: after antibiotic treatment shows bacterial levels and E. coli reads. C: Survival of immunocompromised mice pre-treated (green) with antibiotics or untreated (red) after P. aeruginosa infection [n=10]. D: PA14 CFUs in feces of immunocompromised mice treated with antibiotics (blue) and untreated (red) [**=p<0.005,***=p<0.0005] [n=7-10].

Antibiotic-treated mice subjected to neutropenia via cyclophosphamide injections and infected with *P. aeruginosa* exhibit systemic spread of bacteria (Figure S2). Importantly, all neutropenic dysbiotic mice die within 9 days of oral infection with *P. aeruginosa*. The catalytic role of healthy microbiota is evident by the 90% survival of *P. aeruginosa* infected immuno-compromised mice that are not treated with antibiotics (Figure 2C). Similarly, *P. aeruginosa* load in the stools of infected mice bearing the healthy microbiota are much less than in mice treated with antibiotics, suggesting that commensal microbes inhibit colonisation by *P. aeruginosa* (Figure 2D). To partly re-establish the mouse microbiome we administered a fecal culture supplement (FC) prepared from a pelleted anaerobic stool culture. FC contained the endogenous *Bacteroides, Bifidobacteria* and *Lactobacillus* species and to a lesser extent the endogenous *E. coli* (Figure 3H). The addition of FC in the drinking water had had little to no effect in protecting mice against lethality. However, FC fortified with the endogenous *E. coli* strain (*E. coli* CD1) rescued 70% of mice (Figure 2E). On the other hand, the *E. coli* CD1 in the absence of FC did not protect mice against *P. aeruginosa* infection, (Figure 2E, 2F), suggesting a synergism between the endogenous *E. coli* CD1 and other members of the microbiota as a result of adaptation or co-evolution. Unlike the *E. coli* CD1 strain, the laboratory *E. coli* strain BW25113 showed only a trend in improving mouse survival from *P. aeruginosa* infection, and this effect was not modifiable by FC (Figure 2E, 2F). Despite the marginal effect on survival, *E. coli* BW25113 can stably colonize the mouse gut (Figure S2D) and reduces *P. aeruginosa* burden significantly in the mouse gut within a week post infection (Figure 2G).

**Figure 3:**
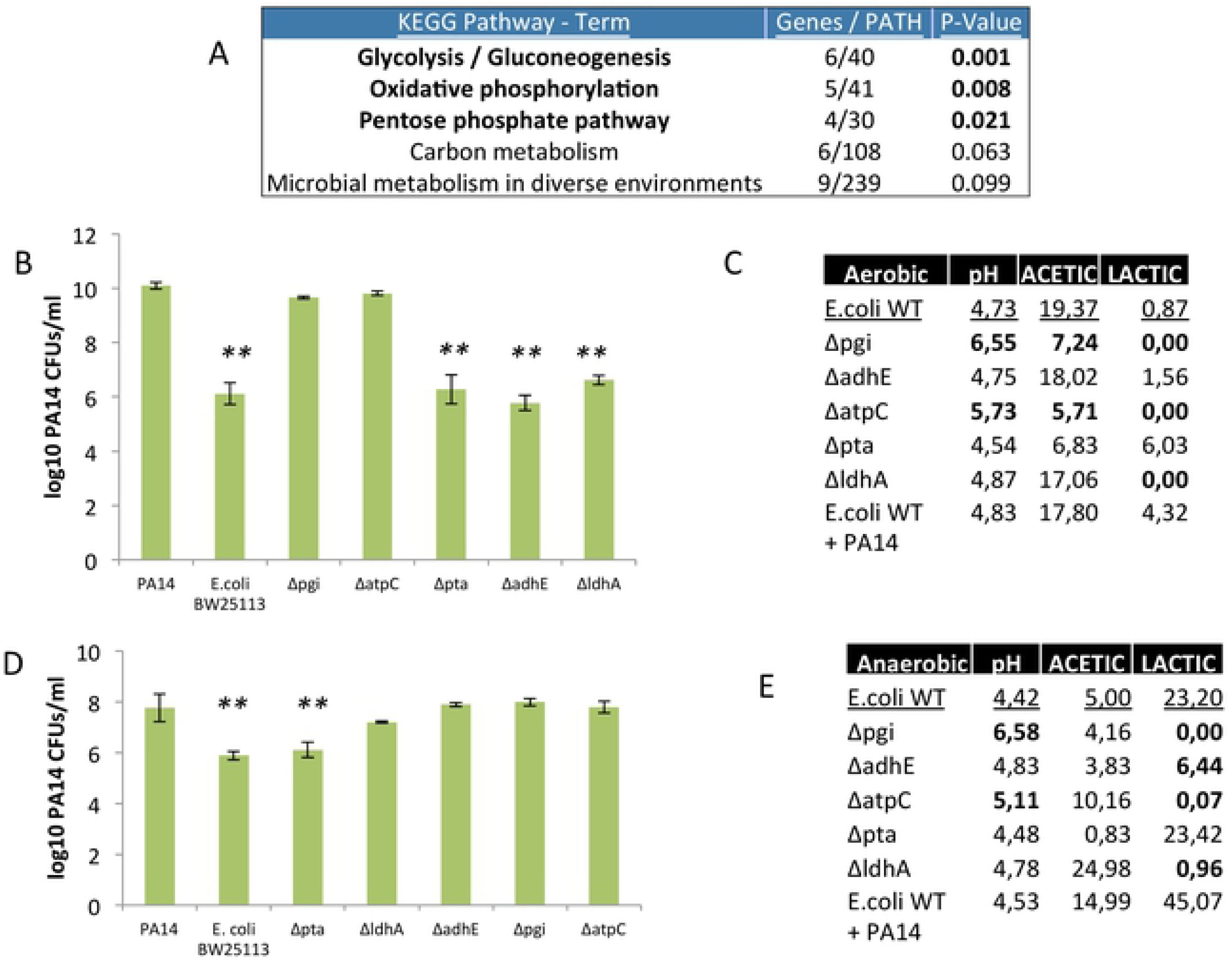
E. coli genes identified to be deficient in inhibiting PA14 and their Acetic and Lactic acid production profiles. A: Gene enrichment analysis identified pathways to be significantly involved in our genome-wide screen results [EASE Score, a modified Fisher Exact P-Value]. B: PA14 CFUs in co-cultures with BW25113 (WT) and isogenic gene mutants identified during aerobic growth [**=p<0.005] [n=3]. C: Liquid culture media pH and acid measurements (mM) 5 hours after bacterial inoculation [n=3]. Bold values are changes observed in our deficient mutants. D: PA14 CFUs in co-cultures with BW25113 (WT) and isogenic gene mutants identified during anaerobic growth [n=3]. E: Liquid culture media pH and acid measurements (mM) 24 hours after bacterial inoculation [n=3]. Bold values are changes observed in our deficient mutants.

### Glucose metabolism pathways in *E. coli* mediate *P. aeruginosa* growth inhibition

*E. coli* QS signalling and the production of the metabolite indole are reported to inhibit *P. aeruginosa* growth (Lee et al. 2009, Chu et al. 2012). To reveal *E. coli* factors that inhibit *P. aeruginosa* in our experiments, we assessed *E. coli* QS mutants and indole production genes previously implicated in bacterial competition (Chu et al. 2012). We found that *E. coli* QS genes luxS and sdiA are not contributing in *P. aeruginosa* inhibition in an LB culture supplemented with 4% glucose (Fig. S3A). In addition, a deletion of the indole production enzyme tryptophanase (*tna*) essentially eliminated indole production, but not the ability of *E. coli* BW25113 to inhibit *P. aeruginosa* (Fig. S3B, S3C). Therefore we performed an unbiased screen of the KEIO collection of 3985 isogenic K-12 BW25113 gene mutants identifying 45 genes that are necessary for the inhibition of *P. aeruginosa* in LB broth supplemented with 4% glucose. Gene enrichment analysis pinpointed glycolysis and the downstream pathways of oxidative phosphorylation and pentose phosphate as strongly enriched (Figure 3A).

To assess the impact of *E. coli* glycolysis and oxidative phosphorylation on the *P. aeruginosa* growth we co-cultured *in vitro P. aeruginosa* with the core glycolysis and oxidative phosphorylation pathway mutants *E. coli, Δpgi* and *ΔatpC*, respectively. In aerobic cultures, wild type *E. coli* BW25113 reduced *P. aeruginosa* concentration by >1,000 fold, while *Δpgi* and *ΔatpC* mutants were unable to inhibit *P. aeruginosa* growth significantly (Figure 3B). This is in line with the fermentation efficiency of the *Δpgi* and *ΔatpC* strains, which was severely compromised with no lactic acid and reduced acetic acid (>2 fold decrease) production and deficient acidification (pH>5.5) of the liquid bacterial culture (Figure 3C). In aerobic conditions lactic acid production is very low compared to acetic acid produced, but none of the mixed acid fermentation mutants, *Δpta, ΔadhE* or *ΔldhA* could abolish production of lactic acid and reduce acetic acid production (Figure 3C). Accordingly, these mutants inhibited *P. aeruginosa* aerobically (Figure 3B). On the other hand, the *Δpgi* and *ΔatpC* strains abolish lactic acid and reduce acetic acid production, and these mutants are the only ones unable to inhibit *P. aeruginosa* (Figure 3B, 3C).

Because the environment in the mammalian gut is anaerobic and the fermentation process towards lactic acid production is much more efficient we further tested this pathway anaerobically. Similarly to aerobic conditions, the core metabolism *E. coli* gene mutants *Δpgi* and *ΔatpC* were unable to inhibit *P. aeruginosa* growth, acidify culture media and produce lactic acid in anaerobic cultures (Figure 3D, 3E). Also *E. coli ΔldhA* and *ΔadhE* mutants exhibited significantly reduced lactic acid production [P<0.001] (Figure 3E) and an impaired ability to inhibit *P. aeruginosa* in an anaerobic culture (Figure 3D). Thus lactic acid production is crucial, while acetic acid production is helpful, in inhibiting *P. aeruginosa* growth.

### Lactic acid and Acetic acid can inhibit *P. aeruginosa* growth and virulence

Supplementation of the *E. coli* mixed acid fermentation by-products acetic acid and lactic acid have been reported to act as antimicrobials against *P. aeruginosa* (Phillips et al. 1968, Levison 1973, Alakomi et al. 2000). We validated the role of these two metabolites in inhibiting *P. aeruginosa* growth at pH 5. Acidic pH of <5 is observed in an *E. coli* culture in the presence of sugars in either aerobic or anaerobic conditions (Figure 3C, 3E). A concentration of 10mM or more of acetic acid, which can be produced by *E. coli* in an aerobic liquid culture (Figure 4C), can totally inhibit *P. aeruginosa* growth (Figure 4A). 10mM or more of lactic acid, which can be produced by *E. coli* in an anaerobic culture (Figure 4D), inhibits *P. aeruginosa* growth (Figure 4B).

**Figure 4.**
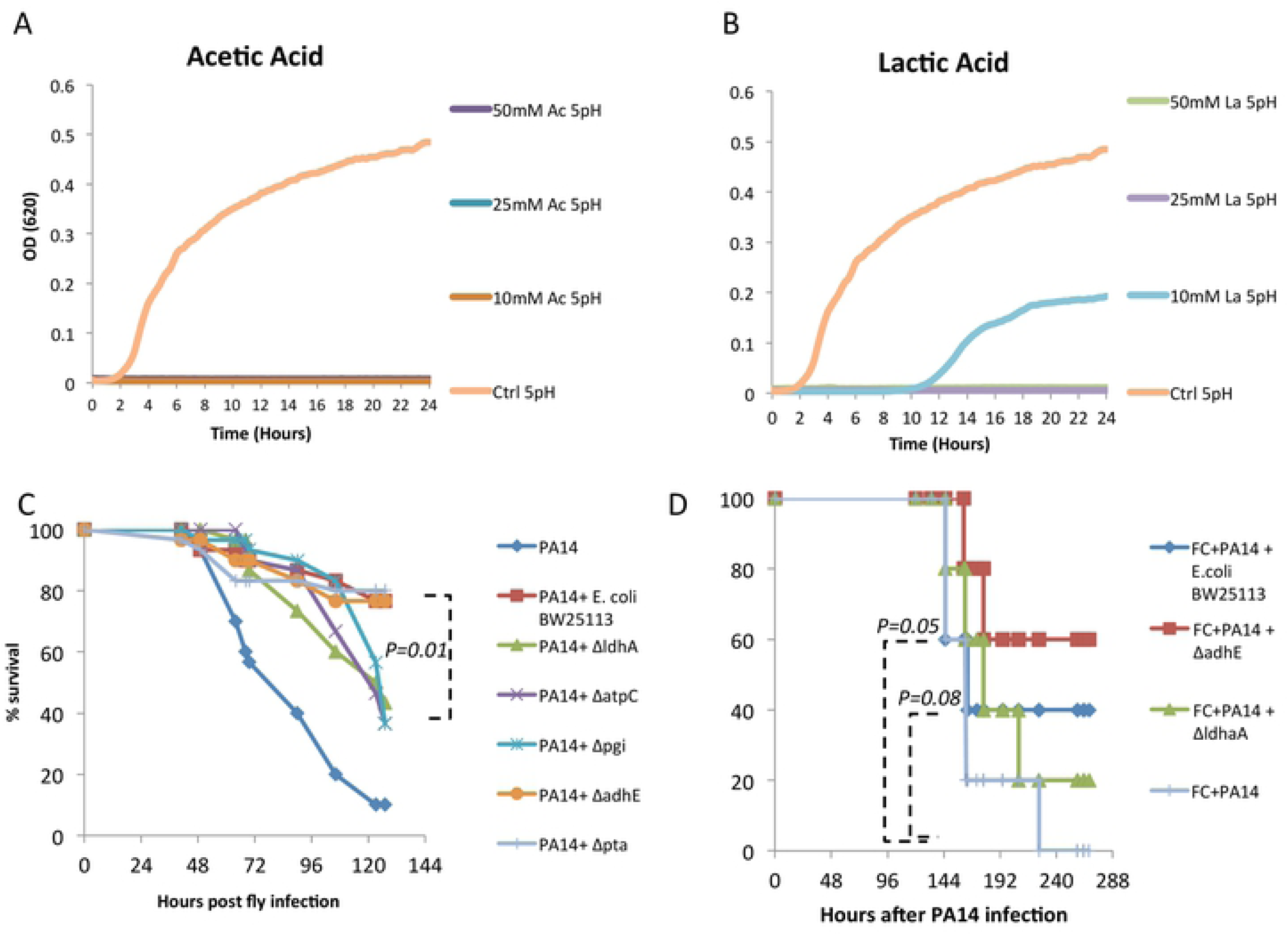
Lactic and Acetic acid capacity to inhibit PA14 growth and virulence in culture media and in the host. A: PA14 growth in LB broth with added Lactic acid concentrations of 10-50 mM at pH 4.5, 5, 7 (Ctrl = 0mM) [n=4]. B: PA14 growth in LB broth with added Acetic acid concentrations of 10-50 mM at pH 4.5, 5, 7 (Ctrl = 0mM) [n=4]. C: Drosophila melanogaster Oregon R survival curve after co-infection with PA14 and BW25113 gene mutants [n=45]. D: Survival of PA14 infected immunocompromised mice complemented with a stool culture and the E. coli strain BW25113 (dark blue), ΔldhA (green), ΔadhE (red). Control antibiotic treated and PA14 infected with no E. coli (light blue), control with no E. coli and no antibiotics (pink) [n=10].

The inhibitory effect of *E. coli* lactic and acetic acid genes was further tested in animal models. In fly infection experiments, the *ΔldhA E. coli* mutant, specifically deficient in lactic acid production, and the core pathway mutants *Δpgi* and Δ*atpC* unable to produce lactic acid anaerobically exhibit diminished ability to rescue flies infected with *P. aeruginosa* (Figure 4C). To the contrary, the Δ*pta* and Δa*dhE* mutants that cannot abolish lactic acid production in culture rescue flies to the levels of the wild type isogenic *E. coli* strain BW25113 (Figure 4C). The same pattern was observed during co-infections in mice. We noticed that the *E. coli* mutant *ΔadhE* significantly rescued 60% of mice from lethality upon oral *Pseudomonas* infection in mice, comparable to the wild type isogenic *E. coli* strain BW25113 rescuing 40% of mice, while the lactic acid defective Δ*ldhA* did not provide any significant rescue (Figure 4D), suggesting that lactic acid is key in inhibiting *P. aeruginosa* in the host.

### In a low-sugar environment *P. aeruginosa* antagonises *Escherichia coli* growth by the unrestricted production of pyocyanin

In the absence of added sugars in the culture media the interaction between *E. coli* and *P. aeruginosa* has an additional level of complexity with *P. aeruginosa* exhibiting the ability to inhibit *E. coli* growth (Figure 5A, 5B). Screening for *P. aeugurinosa* mutants implicated in this process we identified the phenazine system and its regulators to be necessary for *E. coli* growth ihibition by >100 fold in culture (Figure 5A, 5B). Pyocyanin, a redox-active secondary metabolite and a potent antibacterial is produced and secreted by *P. aeruginosa* upon quorum sensing (QS) activation of the phenazine operon (Figure 5C). Supplementation of 10mM of pure pyocyanin was sufficient to inhibit *E. coli* in culture to the same extend as in co-culture with *P. aeruginosa* (Figure 5D, 5A).

**Figure 5:**
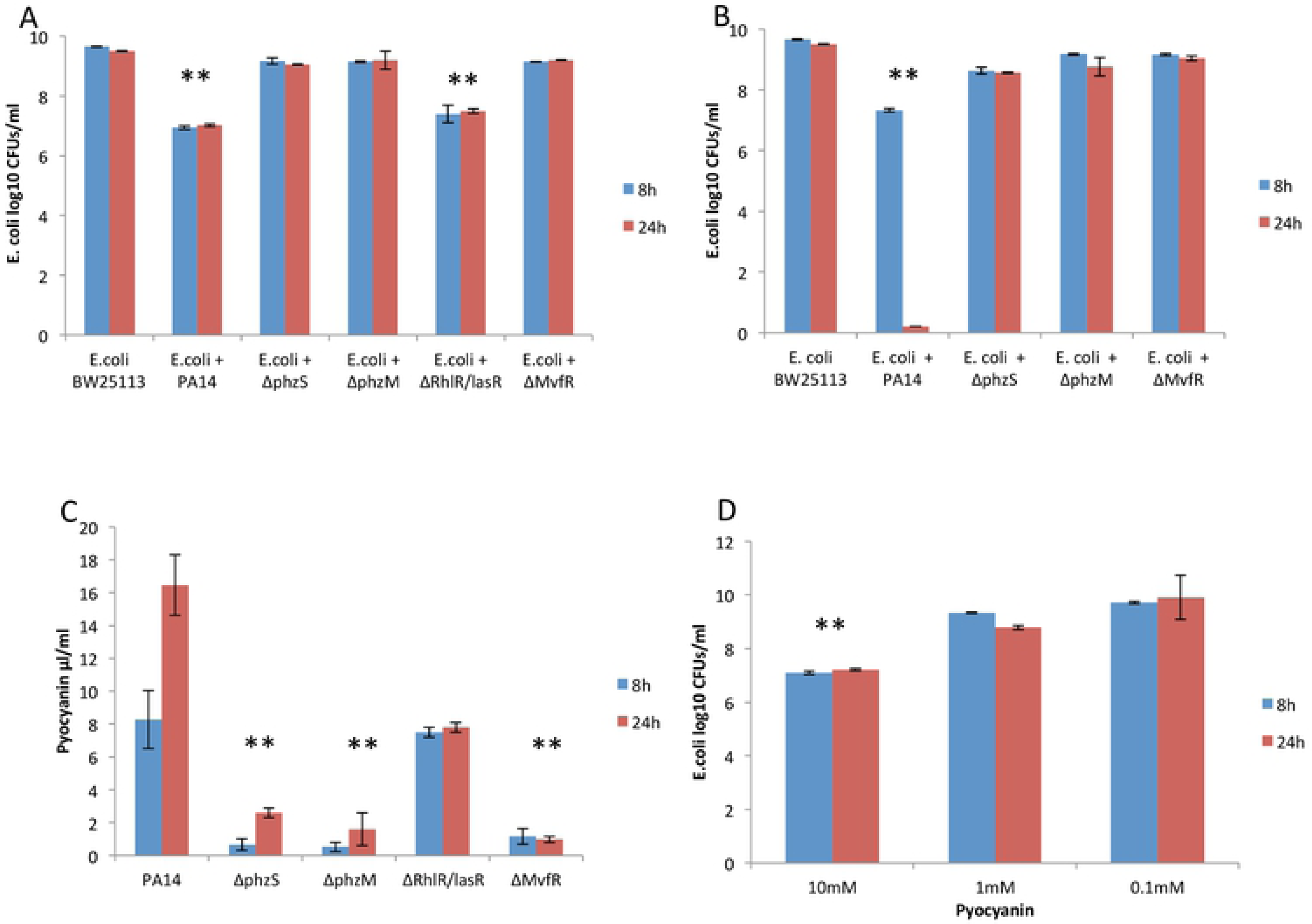
P. aeruginosa toxin pyocyanin inhibits E. coli growth in vitro. A: CFUs of BW25113 growth in co-culture with WT PA14 and QS mutants at 8 (red) & 24 (blue) hours[**=p<0.005] [n=3]. B: CFUs of BW25113 growth in 1:1 LB and supernatant of WT PA14 and QS mutants at 8 & 24 hours [n=3]. C: Pyocyanin concentration (μl/ml) of WT PA14 and QS mutants at 8 & 24 hours [n=3]. D: CFUs of BW25113 growth at 8 & 24 hours in LB with supplemented pyocyanin (mmol/L) [n=3].

Previous work shows that sugars may inhibit the quorum sensing system of *P. aeruginosa* (Wang et al. 2012). We assessed increasing concentrations of glucose and sucrose on PA14 cultures for the QS product pyocyanin and observed that both are able to inhibit pyocyanin production (Figure 6A, 6B). Glucose was tendatively more potent in inhibiting pyocyanin production than sucrose with more than 50% inhibition at the 4% concentration in 24 hours.

**Figure 6:**
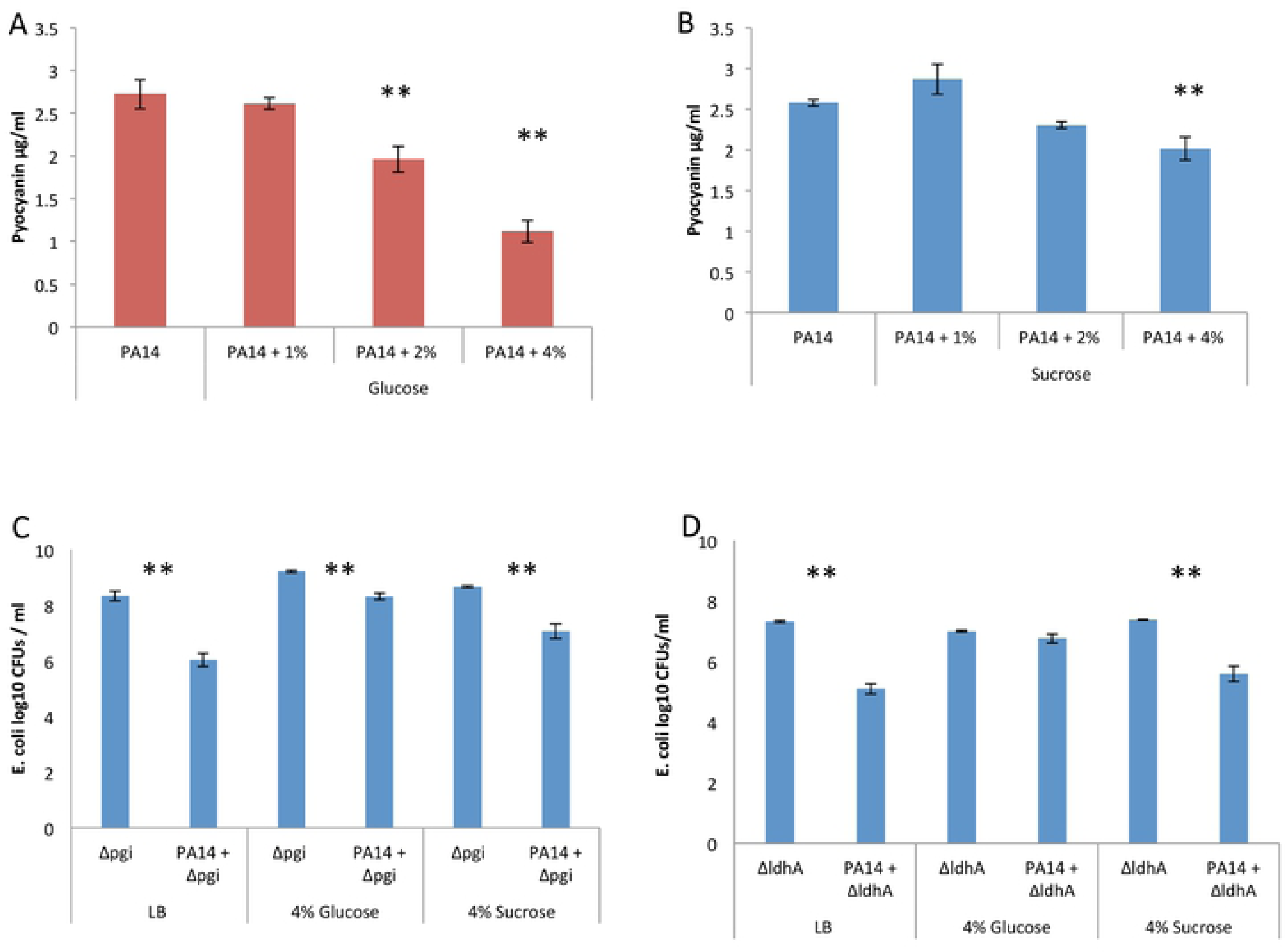
Sucrose and Glucose capacity to inhibit PA14 production of pyocyanin and E. coli inhibition. A: Pyocyanin production (μl/ml) in PA14 LB cultures with added glucose at 24 hours [n=3]. B: Pyocyanin production (μl/ml) in PA14 LB cultures with added sucrose at 24 hours [n=3]. C: E. coli BW25113 Δpgi core glycolysis mutant CFUs at 24 hours in single or co-culture with PA14 [n=3]. D: E. coli BW25113 Δldha lactate dehydrogenase mutant CFUs at 24 hours in single or co-culture with PA14 [n=3]. Plain Luria broth (LB) or with added % of glucose or sucrose was used as culture growth media.

To assess the ability of sugars to interfere with *P. aeruginosa* ability to inhibit *E. coli* we used the *E.coli* mutants *Δpgi* and *Δldha*, which are deficient in glycolysis and lactic acid production, respectively, and are thus unable to inhibit *P. aeruginosa*. We noticed that the addition of 4% glucose reduced the ability of *P. aeruginosa* PA14 to inhibit the gorwth of *E. coli* mutants *Δpgi* and *Δldha* by 1.4-2 log_10_fold (25-100 fold), whereas 4% sucrose only marginally so i.e. by 0.4-0.7 log_10_ (3-5 fold) (Figure 6C, 6D).

Furthermore, we assessed the ability of *P. aeruginosa* to inhibit *E. coli* mutants *Δpgi* and *Δldha* colonization in flies orally infected and fed with 2% or 4% glucose or sucrose. Sugar is a necessary ingredient in the fly food and the sole nutrient in this assay. *Drosophila* colonisation at 0, 2 and 5 days post co-inoculation with the *E.coli* glycolysis mutant *Δpgi* and *P. aeruginosa* showed that, when unable to inhibit *P. aeruginosa, E. coli* is inhibited by *P. aeruginosa* at 2 and 5 days in the fly gut (Figure 7A, 7B). Moreover, *P. aeruginosa* can inhibit the colonization of *E.coli* lactic acid mutant *Δldha* immediately upon co-inoculation at day 0 (Figure 7C). These results suggest that, 2-4% of dietary sugars do not affect *P. aeruginosa* growth and ability to inhibit gut colonization by *E. coli* mutants unable to ferment sugars into lactic acid.

**Figure 7:**
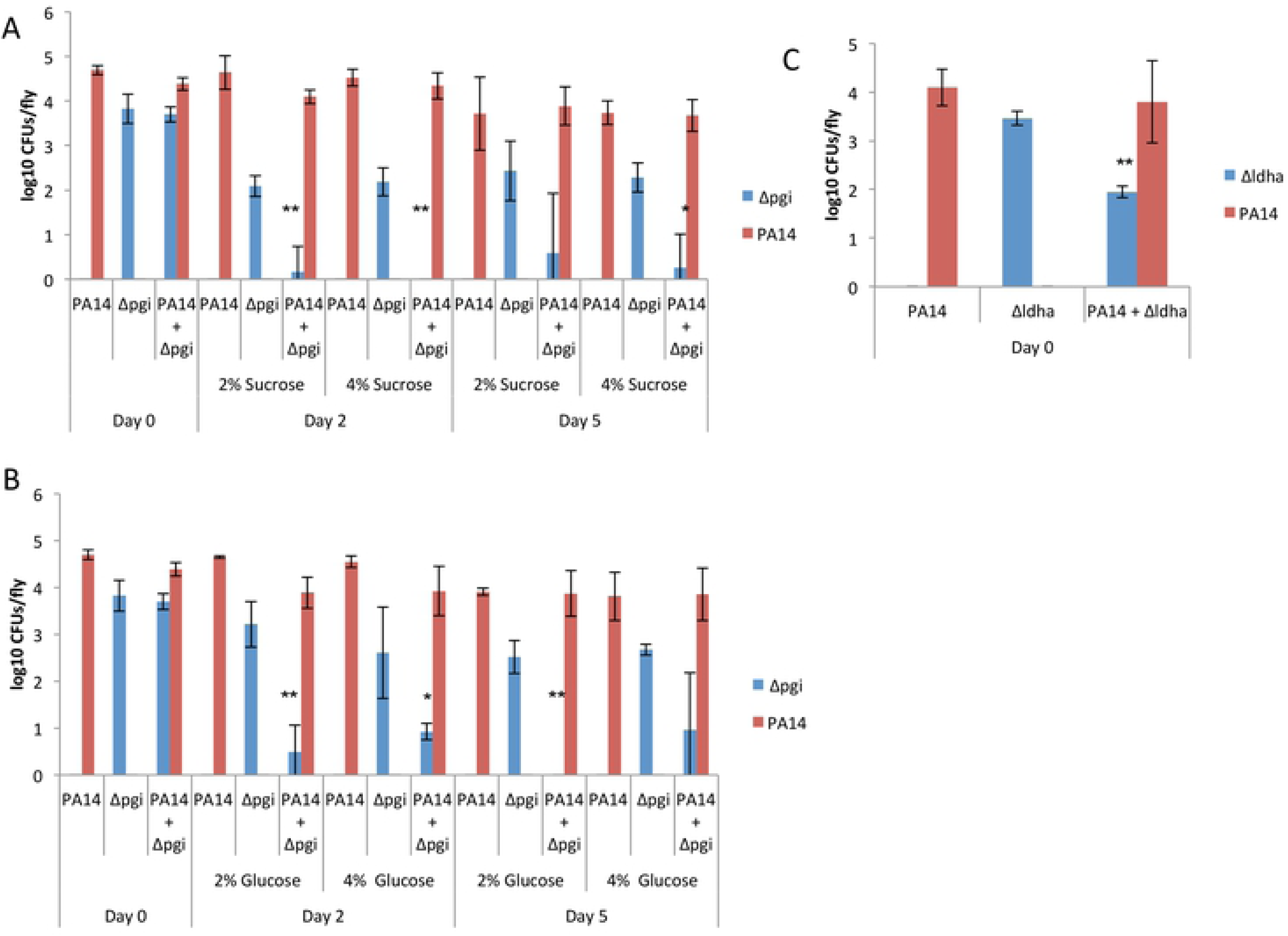
Sucrose and Glucose capacity to inhibit PA14 production of pyocyanin and E. coli inhibition. A: E. coli BW25113 Δpgi core glycolysis mutant (blue) and PA14 (red) colonisation CFUs at 2 and 5 days after fly infection. 2 % and 4% sucrose was used in infection media and as fly food [n=3]. B: E. coli BW25113 Δpgi core glycolysis mutant and PA14 colonisation CFUs at 2 and 5 days after fly infection [n=3]. 2 % and 4% glucose was used in infection media and as fly food [*=p<0.05,**=p<0.005]. C: E. coli BW25113 ΔldhA lactate dehydrogenase mutant and PA14 colonisation CFUs at day 0 after fly infection [n=3].

### Diet contribution in the interaction of *E. coli* and *P. aeruginosa* in the mouse gut

Diet is very important for the maintenance of a healthy microbiome and in shaping the intestinal immune response (Brown et al. 2012, Hooper et al. 2012, Tilg and Moschen 2015). Our study shows that the interaction between *E. coli* and *P. aeruginosa* is shaped by the presence of high concentrations of sugars. Therefore, we sought to investigate in mice the contribution of three nutritionally extreme diets: a protein-based, a fat-based and a carbohydrate-based. In the carbohydrate-based diet the total levels of sugars (sucrose, glucose and fructose) in the feces were the 67.4μg/ml, which was higher than any of the other diets, while only 18.7 μg/ml in the presence of orally administered *E. coli* (Figure S4A). This means that *E. coli* is consuming sugars in mouse gut. Yet, lactic acid levels were the highest in the fat-based diet in the presence of *E. coli* (Figure S4B). Accordingly, *P. aeruginosa* CFUs were reduced by *E. coli* in mice fed with the fat-based diet (Figure 8B), but not with the carbohydrate or the protein-based diets (Figure 8A, 8C). Similarly, mouse survival upon *P. aeruginosa* infection in neutropenic mice was the highest in mice fed with fat-based diet and co-inoculated with *E. coli* as opposed to mice fed with the carbohydrate-based diet and co-inoculated with *E. coli* (Figure 8D). However, the fat-based diet does not favour *E. coli* gut colonization, as the *E. coli* CFUs in the feces are comparable between the carbohydrate and the fat-based diets and lower than those of the protein-based diet (Figure 8E).

**Figure 8:**
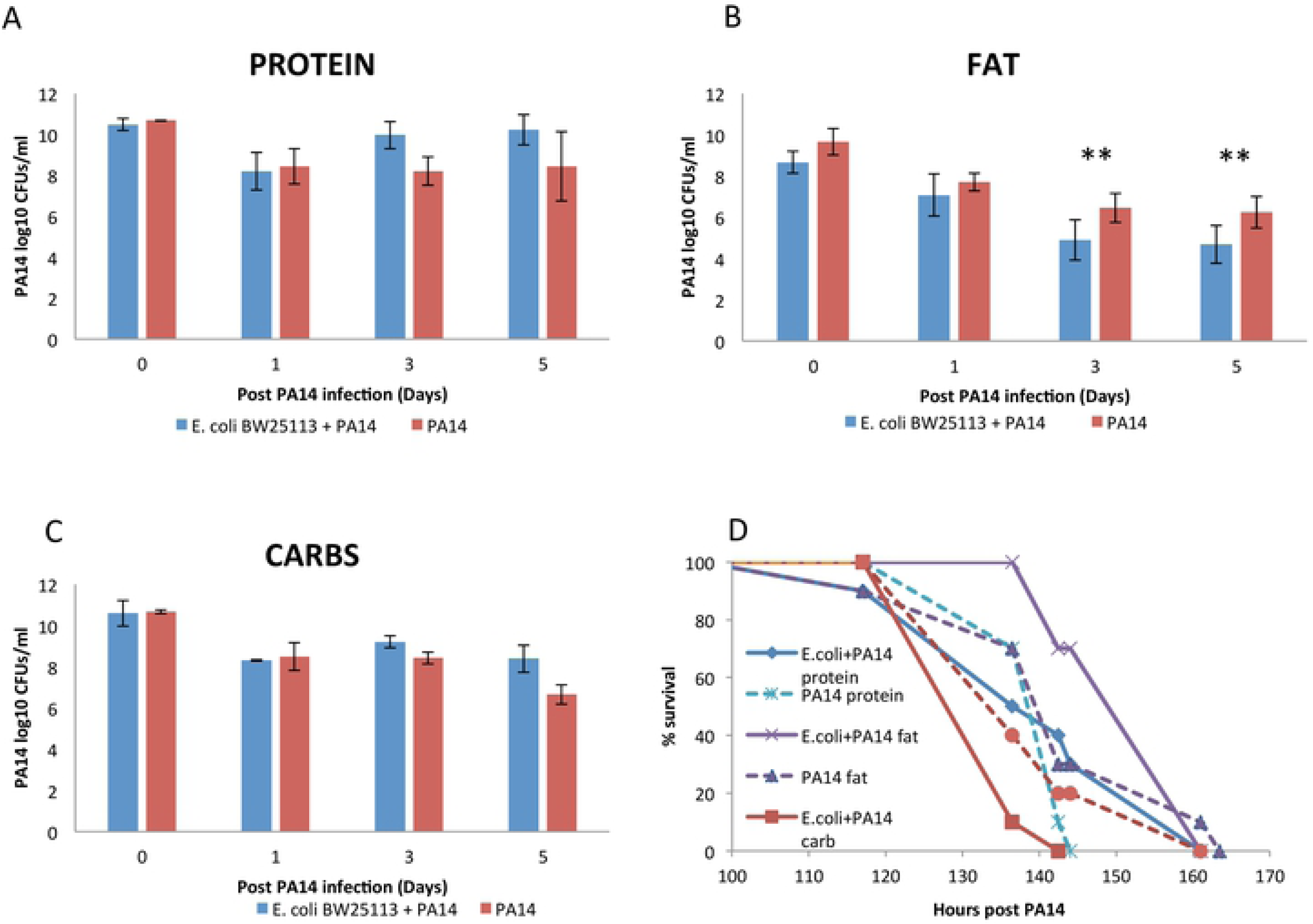
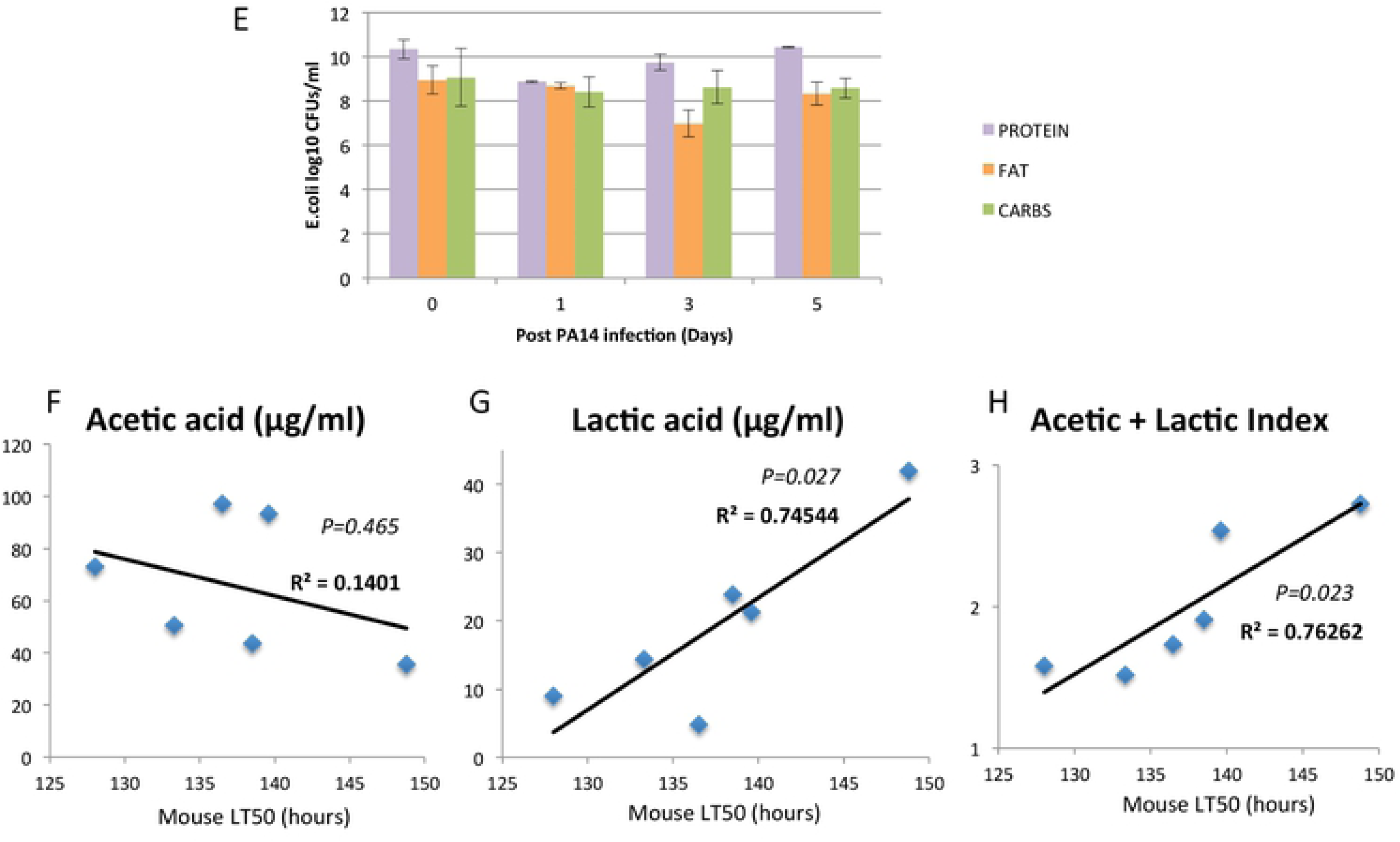
Diet effects on P. aeruginosa mouse colonisation and interaction with E. coli. A: PA14 CFUs in feces of immunocompromised mice fed on high protein diet and infected with PA14 (red) and PA14 + E. coli (blue) [n=10]. B: PA14 CFUs in feces of immunocompromised mice fed on high fat diet and infected with PA14 (red) and PA14 + E. coli (blue) [n=10]. C: PA14 CFUs in feces of immunocompromised mice fed on high carbohydrate diet and infected with PA14 (red) and PA14 + E. coli (blue) [**=p<0.005] [n=10]. D: Survival of PA14 infected immunocompromised mice complemented with E. coli strain BW25113 (blue, purple, red) fed on high fat, carb and fat diets. [p-value= Fat PA14+E. coli vs Protein PA14+E.coli p=0.04, Fat PA14+E.coli vs Carb PA14+E.coli p=0.0001, Protein PA14+E.coli vs Carb PA14+E.coli p=0.06] [n=10]. E: E. coli CFUs in feces of immunocompromised mice fed on high protein (purple), high fat (orange) and high carbohydrate diet (green) infected with PA14 and E. coli [n=10]. F and G: Pearson correlation of Acetic and Lactic acid concentration for each of the 6 mouse diets in the mouse feces with the average time of death (LT50) of the corresponding mice. H: A combinatorial index of normalized mouse fecal Acetic and Lactic acid concentration correlated withLT50.

To assess if fecal acetic and lactic acid production may indicate protection against *P. aeruginosa*, we correlated acid concentration in the feces with the lethal time 50% (LT50) of the corresponding sets of mice. We found that lactic but not acetic acid level alone correlates significantly and positively with survival to infection (Figure 8F, 8G), while an index of normalized concentration values for acetic and lactic acid combined gave also a clear correlation with survival (Figure 8H). We conclude that the standard balanced diet (Figure 2G) and the fat-based diet (Figure 8B), rather than the carbohydrate-based diet (Figure 8C) facilitate the inhibitory effect of *E. coli* on *P. aeruginosa*; and we suggest that, given the complexity of the mammalian intestinal environment, the metabolic output in acetic and lactic acid production rather than the dietary input is indicative of susceptibility to intestinal *P. aeruginosa* infection.

## Discussion

Despite bacterial antagonism being observed since 1905 (Rettger 1905), there are not many cases supporting the alternative model of one pathogen one colonization resistor, according to which specific bacterial strains protect the host against infection (Byrd and Segre 2016). Known cases include the inhibitory effect of *Clostridium scindens* in resistance against *C. difficile* infection (Buffie et al. 2015), of the non-toxigenic against the enterotoxigenic *Bacteroides fragilis* (Wagner et al. 2016), of *E. coli* O21:H+ against muscle atrophy due to infection, and of *E. coli* EcN against intestinal pathogens (Byrd and Segre 2016). Antagonistic interactions between a pathogenic and a non-pathogenic bacterial strain may include: (a) the direct inhibition of pathogen’s growth, colonization or virulence by the non-pathogenic strain, or (b) the indirect effect of the non-pathogenic strain in inducing or supporting the host defense to infection. To establish the mode of interaction between *E. coli* and *P. aeruginosa* we first examined whether *Drosophila* can sustain the colonization of these species in the fly. Sugar-based diets allow stable bacterial colonization with either *P. aeruginosa* or *E. coli* strains. When flies are co-infected with both species, *P. aeruginosa* is significantly reduced or eliminated. In addition, only *P. aeruginosa* can be highly lethal to the flies and the presence of *E. coli* dramatically reduces lethality. Antagonism between *P. aeruginosa* and other species is nevertheless species specific. Previous studies show that human orophangeal bacteria, which are predominantly gram-positive and differ from those of the human intestine, include *Neisseria, Streptococcus, Staphylococcus* and *Actinomyces* species, and many of them induce rather than antagonize *P. aeruginosa* virulence (Sibley et al. 2008). Moreover, peptidoglycan, which is abundant in gram-positive bacteria, can directly induce the virulence of *P. aeriginosa* (Korgaonkar et al. 2013). Thus, *E. coli,* as opposed to gram-positive lactic acid bacteria, might serve as a safer inhibitor of *P. aeruginosa*, by inhibiting its growth without inducing its virulence.

*P. aeruginosa* primarily affects hospitalised and immunocompromised individuals. It is usually a life-threatening condition during severe burn wound and lung infection, but humans carry *P. aeruginosa* usually asymptomatically in their intestines (Stoodley and Thom 1970). One reason for *P. aeruginosa* being benign in the human gut should be through the action of intestinal microbiota, which are part of the host defense to intestinal infection (Kamada et al. 2013, Schubert et al. 2015, Ge et al. 2017). Previous studies describe the use of antibiotic cocktails that favor *P. aeruginosa* intestinal colonization by compromising resistance by the intestinal microbiota (Hentges et al. 1985). Accordingly, we show that antibiotic use in mice diminishes all the prevalent phyla, eradicates *E. coli,* and induces dysbiosis. Using a *Pseudomonas*-induced gut-derived sepsis model to investigate dissemination and lethality in neutropenic mice (Koh et al. 2005, Matsumoto et al. 2008), we found that mice not-given antibiotics exhibited 90% survival and lower colonization of *P. aeruginosa* compared to the 0% survival of antibiotic-treated animals. In addition, reintroduction of the commensal microbes through a fecal culture of 8 potentially beneficial bacterial species was inefficient in improving mouse protection from lethality. Nevertheless, a high dose of the endogenous *E. coli* CD1 isolate exhibited 70% protection against *P. aeruginosa* infection, but only when supplemented with the fecal culture. We postulate the symbiotic adaptation of the mouse-isolated *E. coli* strain with the mouse gut environment and its microbiota.

Through genome-wide screening in flies we pinpointed *E. coli* glucose metabolism and fermentation mutants deficient in lactic and acetic acid production responsible for inhibiting *P. aeruginosa*. In antibiotic-treated mice, a similar trend was observed where the lactate dehydrogenase deficient *E. coli* mutant was unable to protect mice from *P. aeruginosa* infection and mortality. The anti-infective properties of lactic acid may be attributed to lowering the pH, but also to the permeabilization of the outer membrane of gram-negative bacteria (Alakomi et al. 2000). On the other hand, *P. aeruginosa* produces many virulence factors regulated by QS, such as pyocyanin, which has bactericidal properties (Baron and Rowe 1981). Accordingly, we notice that only strains of *P. aeruginosa* able to produce pyocyanin can inhibit *E. coli*. Interestingly, a high concentration of sugars may inhibit *P. aeruginosa* QS (Wang et al. 2012), and adding glucose or sucrose in *P. aeruginosa* cultures, pyocyanin production is compromised. Thus, sugars not only fuel *E. coli* fermentation towards lactic acid production, but may also inhibit *P. aeruginosa*’s ability to fight *E. coli*.

The role of diet has been extensively studied in response to gut microbiota and host physiology (Singh et al. 2017). Hence, we explored three different diets, based either on carbohydrates (cornstarch and sucrose), fat (vegetable shortening) or protein (casein). Mice feeding on these diets exhibited complex features: First, the carbohydrate-based diet did not improve but rather reduced the ability of *E. coli* to inhibit *P. aeruginosa* colonization and concomitant mortality. This might be because this carbohydrate-based diet does not deliver a significant amount of free sugars in the mouse colon. While sugars are higher in the feces of mice fed with a carbohydrate-based diet, they may be quite low to have the anticipated impact on *E. coli*. Second, the protein-based diet sustains more *E. coli* than the other diets, yet, this didn’t translate into better inhibiting capacity against *P. aeruginosa*. This might be because casein inhibits or lacks the ability to fuel fermentation into lactic acid. Third, the fat-based diet without sustaining more *E. coli* compared to the other diets, allows *E. coli* to produce more lactic acid that can inhibit *P. aeruginosa* growth. This is in line with evidence showing that unsaturated fat may benefit lactic acid bacteria in mice (Caesar et al. 2015).

Lactic acid in the mouse feces is much lower than the lowest inhibitory concentration tested in culture. Nevertheless, *E. coli* mutants defective in lactic acid production are also defective in inhibiting *P. aeruginosa* in the fly and mouse gut. The ability of any chemical to inhibit bacterial growth depends on the environment and thus additional factors (e.g. additional antimicrobials) in the fly and mouse gut may boost the ability of lactic acid to inhibit *P. aeruginosa*. In addition, high sugar levels may be difficult to achieve by a high carbohydrate diet because sugar is readily absorbed in the small intestine, and *E. coli* and other commensals may use dietary fat more efficiently towards lactic acid production. Thus, the metabolic output in the colon rather than the dietary input might better dictate the balance between and among bacterial species. Metabolic output nevertheless is a result of diet, microbiota composition and the host physiology acting in concert. Accordingly, fecal metabolomics might prove very helpful in predicting the outcome of bacterial interactions in the human colon.

## Acknowledgements

We thank Christos Shammas from AVVA Pharmaceuticals for 16S sequencing. James Angus Chandler and Robert J. C. McLean for Lactobacillus and *E. coli* strains and the KEIO collection (Japan) for the BW25113 library strains. Tassos Economou for the Enteropathogenic *E. coli* O127:H6 E2348/69 strain. Laurence Rahme for critical reading and editing. We also thank Marie Curie CIG and Fondation Sante for funding to YA.

## Supporting information file Legends

**Figure S1: E. coli inhibits P. aeruginosa growth and virulence in the Drosophila gut and in culture in the presence of glucose.** A: Survival of Drosophila melanogaster Oregon R flies infected with P. aeruginosa strain PA14 (green), E. coli strain BW25113 (yellow) or co-infected with E. coli and PA14 (red) [n=45]. B: Colonisation levels measured in colony forming units (CFUs) at Day 2 or Day 5 post PA14 infection (green), BW25113 (yellow) in mono or co-infected flies [n=3]. C: CFUs of PA14 growth in the presents or absence of 4% glucose or fructose and E. coli BW25113 in Luria-Bertani cultures [n=3]. D: Optical density measurements at 600nm of PA14 growth in liquid supernatant of E. coli cultures +/- 4% glucose [n=9].

**Figure S2.** A, Internal organs of an untreated mouse. B, Internal organs of an antibiotic treated and infected mouse. C, Log10 CFUs per ml of blood or whole organ counts in Spleen, liver, lung and mesenteric lymph nodes. D, E. coli BW25113 colonisation levels in mice infected for 7 Days with PA14 followed by 1 Day with E. coli at a concentration of 3×10^8 bacteria in the drinking water. CFU measurements in fecal samples 1, 3, 5 and 7 days after PA14 infection.

**Figure S3: E. coli quorum sensing and Indole against P. aeruginosa growth.**

A: PA14 CFUs at 24 hours when co-cultured with wild type BW25113 or quorum sensing E. coli mutants sdiA and luxS [n=3].

B: Indole measurements in wild type E.coli strain BW25113 and tryptophanase Δtna mutant cultures at 6 (blue) and 24 (red) hours [n=3].

C: PA14 CFUs at 24 hours in co-culture with wild type BW25113 and tryptophanase Δtna mutant. [**=p<0.005], [n=3].

**Figure S4: Sugar and acid concentrations of mouse feces reared on nutrient-defined diets.** A: Sugar measurements in samples of mice feces reared on each diet (μg/ml). B: Lactic and acetic acid concentrations measurements in samples of mice feces reared on each diet (μg/ml).

